# Selective REM-Sleep Suppression Increases Next-Day Negative Affect and Amygdala Responses to Social Exclusion

**DOI:** 10.1101/2020.06.15.148759

**Authors:** Robert W Glosemeyer, Susanne Diekelmann, Werner Cassel, Karl Kesper, Ulrich Koehler, Stefan Westermann, Armin Steffen, Stefan Borgwardt, Ines Wilhelm, Laura Müller-Pinzler, Frieder M Paulus, Sören Krach, David S Stolz

## Abstract

Healthy sleep, positive general affect, and the ability to regulate emotional experiences are fundamental for well-being. In contrast, various mental disorders are associated with altered rapid eye movement (REM) sleep, negative affect, and diminished emotion regulation abilities. However, the neural processes mediating the relationship between these different phenomena are still not fully understood. In the present study of 42 healthy volunteers, we investigated the effects of selective REM sleep suppression (REMS) on general affect, as well as on feelings of social exclusion, emotion regulation, and their neural underpinnings. Using functional magnetic resonance imaging we show that REMS increases amygdala responses to experimental social exclusion, as well as negative affect on the morning following sleep deprivation. There was no evidence that emotional responses to experimentally induced social exclusion or their regulation using cognitive reappraisal were impacted by diminished REM sleep. Our findings indicate that general affect and amygdala activity depend on REM sleep, while specific emotional experiences possibly rely on additional psychological processes and neural systems that are less readily influenced by REMS.

## Introduction

Sleep is fundamental for general well-being ^[1,2]^. Healthy sleep consists of two repeatedly cycling types of sleep: non-rapid eye movement sleep (NREM) including slow-wave sleep (SWS) phases of deep sleep and rapid eye movement sleep (REM). Over the course of the night, the percentage of SWS decreases while REM sleep percentage increases towards the morning ^[3]^. In many occasions however, sleep is fragmented or shortened (e.g. due to stress, early school or work starts and late bedtimes) and even minor sleep deprivation can have broad, short-term as well as long-term consequences for health and well-being. Sleep disturbances affect health on the level of immune regulation ^[4–6]^ and metabolic markers ^[7,8]^. Furthermore, sleep loss negatively impacts cognitive processing and emotional reactivity ^[9,10]^, as well as social behavior, leading to withdrawal from social interaction and feelings of loneliness [11].

Interestingly, most mental disorders are associated with sleep peculiarities. Insomnia is comorbid in 85-90% of individuals diagnosed with major depression ^[12–14]^. Alterations in REM sleep architecture (REM sleep latency, density and distribution) are particularly prominent in most mental disorders such as major depression, bipolar disorder and Posttraumatic Stress Disorder (PTSD) ^[15,16]^. Given that these disorders are mainly characterized by altered affective experience during daytime, it is interesting to note that inadequate sleep itself has been shown to impact next day emotionality. Sleep loss has been shown to reduce the ability to regulate emotions, leading to exaggerated responses to aversive or even neutral experiences ^[17,18]^. Recent studies even suggest sleep abnormalities as being causal agents in mood disorders rather than just a symptom ^[13,19]^.

Although single sleep stages seem to be differentially involved in mood disorders, the role of these sleep stages for cognitive and emotional functioning is not well understood. Increasing evidence suggests that SWS and associated neurophysiological processes support the consolidation of newly encoded declarative memories ^[20,21]^. REM sleep, on the other hand, is particularly important for emotional processing, including emotional reactivity ^[22,23]^ and the formation of emotional memories ^[24,25]^. REM-sleep dreaming was also found to attenuate residual emotional load from the day before ^[26,27]^. However, only very few studies have examined the effects of selective suppression of REM sleep during an otherwise normal night of sleep. The existing evidence suggests that the selective deprivation of REM sleep mainly disturbs the consolidation of emotional memories ^[28–30]^, whereas the selective suppression of SWS mainly impairs emotionally-neutral declarative memory encoding and consolidation ^[31,32]^. These experimental findings are in line with clinical observations suggesting that REM sleep in particular is closely tied with emotional functioning ^[19]^.

In order to better understand the processes underlying the interaction of sleep and emotion, research increasingly addresses the neural mechanisms associated with REM sleep-dependent emotional functioning. At the neural level, a reduction of emotional reactivity in response to previously learned emotional events was found to be accompanied by an overnight decrease of amygdala reactivity, a cluster of nuclei in the temporal lobes and part of the limbic system ^[27,33]^. The limbic system has been widely associated with the processing of affectively laden stimuli and plays an important role in the guidance of behavioral responses to such stimuli ^[34,35]^. Under conditions of total sleep deprivation, next-day negative affect is accompanied by increased amygdala reactivity and decreased functional coupling of the medial prefrontal cortex (MPFC) with limbic structures ^[36]^. Since the MPFC is thought to exert inhibitory control on the amygdala ^[37]^, this finding has been interpreted to reflect a failure of top-down control in the regulation of appropriate emotional responsivity ^[36]^. Simon and colleagues directly examined the effect of sleep deprivation on the neural correlates of next day emotion reactivity ^[17]^. Following sleep deprivation, there was no indication for valence-specific processing of affective pictures in the amygdala, however in the sleep-rested control night, a low amount of REM sleep was associated with a decline in anterior cingulate cortex (ACC)-amygdala connectivity, possibly reflecting a specific effect of REM sleep on cognitive control of emotions.

Successful cognitive control of emotions is regarded to be an essential prerequisite of mental health ^[38,39]^. In daily life, emotions are constantly regulated either implicitly or explicitly by applying specific cognitive strategies like suppression (e.g. distracting the attention away from unpleasant emotional experiences) or reappraisal (e.g. reinterpreting an unpleasant emotional situation). Emotion regulation thus refers to the ability to “influence which emotions we have, when we have them, and how we experience and express these emotions” (p.497 ^[40]^). Interestingly, correlational evidence indicates that the success of emotion regulation is associated with sleep quality ^[41]^. In this study, participants were asked to engage in cognitive reappraisal (CRA) ^[42]^, in this case applying a previously learned cognitive strategy to “redirect the spontaneous flow of emotions” (p. 6 ^[43]^), while watching a sadness-inducing film. The ability to decrease self-reported sadness using CRA compared to baseline was lower in participants who reported poorer sleep quality during the preceding week ^[41]^.

Despite their substantial clinical significance, the neural mechanisms of the effect of REM sleep on the efficacy of regulation strategies in ameliorating unpleasant affect remain unknown. To fill this gap, it is important to experimentally induce negative affect that may then be attenuated by applying CRA strategies. In order to induce negative affect in the present study, we simulated social exclusion in a laboratory setting using the so-called *Cyberball* ^[44]^. *Cyberball* is a virtual ball-tossing paradigm, where participants are playing with a preset computer program while believing that they are playing with two other human participants. By manipulating the number of ball-tosses towards the participant, the degree of social inclusion can be controlled experimentally. Evidence suggests that the distressing experience of social exclusion might share neural regions and mechanisms with the affective processing of physical pain ^[45–47]^. In a neuroimaging study using the *Cyberball* game, participants showed greater activations in cingulate and prefrontal regions involved in the processing of the affective pain component when excluded compared to being included ^[48]^. Although behavioral consequences and neural activation patterns associated with social exclusion have been studied quite intensively, there is scarce research on intervening cognitive appraisals and coping mechanisms regarding feelings of social exclusion ^[49]^.

The aim of the present study was to examine how REM sleep suppression impacts the following day general affect, emotional reactivity, and associated neural mechanisms of emotion regulation during the acute experience of social exclusion. In a between-subjects design we invited participants to a combined polysomnography and fMRI study. After a habituation night allowing regular sleep, for the second, experimental night participants were randomly allocated to either a REM sleep suppression (REMS) group or one of two control groups: a non-suppression control group with regular sleep (CTL) and a high-level control group with similar amounts of awakenings, but where suppression targeted phases of slow wave sleep (SWSS). To assess the impact of REMS on general affect, subjects repeatedly filled in the Positive and Negative Affect Schedule (PANAS) ^[50]^. In the morning after the experimental night, subjects participated in the *Cyberball* during fMRI scanning to induce feelings of social exclusion. All participants engaged in two sessions of the game. In the first session, participants played the game without any instructions. In the second session, participants were instructed to actively regulate their emotions by applying the previously learned CRA.

We hypothesized that selective REMS (vs. SWSS and regular sleep) generally reduces positive and increases negative affect. Furthermore, we expected that selective REMS (vs. SWSS and regular sleep) increases emotional reactivity during social exclusion and dampens the effect of CRA on emotional reactivity. On the level of neural systems we explored whether REMS leads to altered functional activity in (para-)limbic areas such as the amygdala, ACC, insula, and hippocampus during social exclusion and whether neural activity of these regions is modulated by targeted REMS during cognitive reappraisal.

## Results

### Experimental Sleep Manipulation Selectively Reduces REM Sleep Percentage

The experimental sleep manipulation (see figure 1) successfully suppressed REM sleep in the REMS group during the experimental night (REMS score = %REM sleep in habituation – %REM sleep in experimental night; effect of group: *F*(2,37)=13.21, *p*<.001, *η*^2^=.42; planned contrast REMS > others: *t*(38)=5.17, one-sided *p*<.001). The SWSS and CTL groups were similar with regard to REMS (*t*=0.52, two-sided *p*=.608; see table 1 and figure 2 for details).

**Table 1.** Sleep Measures

| n subjects with valid<br>PSG dataset | CTL |  |  |  | REMS |  |  |  | SWSS |  |  |  |
| --- | --- | --- | --- | --- | --- | --- | --- | --- | --- | --- | --- | --- |
|  | Habituation night |  | Experimental night |  | Habituation night |  | Experimental night |  | Habituation night |  | Experimental night |  |
|  | 15 |  | 15 |  | 16 |  | 17 |  | 9 |  | 10 |  |
|  | *M* | *SD* | *M* | *SD* | *M* | *SD* | *M* | *SD* | *M* | *SD* | *M* | *SD* |
| TST (min) | 369.94 | 78.06 | 414.59 | 26.08 | 410.86 | 40.24 | 399.05 | 39.02 | 384.63 | 54.15 | 402.96 | 31.28 |
| N1 (% TST) | 1.87 | 2.76 | 0.58 | 0.96 | 0.60 | 0.65 | 1.72 | 2.54 | 2.47 | 2.57 | 1.18 | 1.39 |
| N2 (% TST) | 61.82 | 7.27 | 61.11 | 8.53 | 64.83 | 4.69 | 68.66 | 7.62 | 65.68 | 8.44 | 69.33 | 8.29 |
| SWS (% TST) | 17.11 | 6.21 | 18.04 | 7.92 | 14.18 | 6.88 | 16.86 | 5.62 | 13.98 | 7.43 | 9.83 | 4.82 |
| REM (% TST) | 19.21 | 6.33 | 20.27 | 4.39 | 20.39 | 5.86 | 12.76 | 6.16 | 17.42 | 6.56 | 19.65 | 8.87 |
| WASO (min) | 67.16 | 50.46 | 51.06 | 29.00 | 47.01 | 40.18 | 71.10 | 42.66 | 57.57 | 38.50 | 61.63 | 37.96 |
| WASO/(TST+WASO) | 0.16 | 0.13 | 0.11 | 0.06 | 0.10 | 0.09 | 0.15 | 0.09 | 0.13 | 0.10 | 0.13 | 0.08 |
| Awakenings | 8.13 | 3.09 | 9.00 | 4.21 | 7.75 | 3.66 | 11.47 | 4.90 | 11.11 | 6.64 | 15.00 | 6.68 |
| REM latency (min) | 142.97 | 68.32 | 106.10 | 51.71 | 101.19 | 43.15 | 154.12 | 93.11 | 137.78 | 86.27 | 123.35 | 82.01 |

Note. In both the REMS group and the SWSS group, PSG data quality from one subject was not suitable for analysis in the habituation night. CTL=Control group with undisturbed sleep; REMS=rapid eye movement sleep suppression group; SWSS=slow wave sleep suppression group; TST=total sleep time (time from sleep onset to morning awakening after the time awake during the night is subtracted); N1=Sleep stage 1, N2=Sleep stage 2, SWS=Sleep stage N3; REM=rapid eye movement sleep; WASO=wakefulness after sleep onset to morning awakening; *M*=Mean; *SD*=Standard Deviation

**Figure 1.**
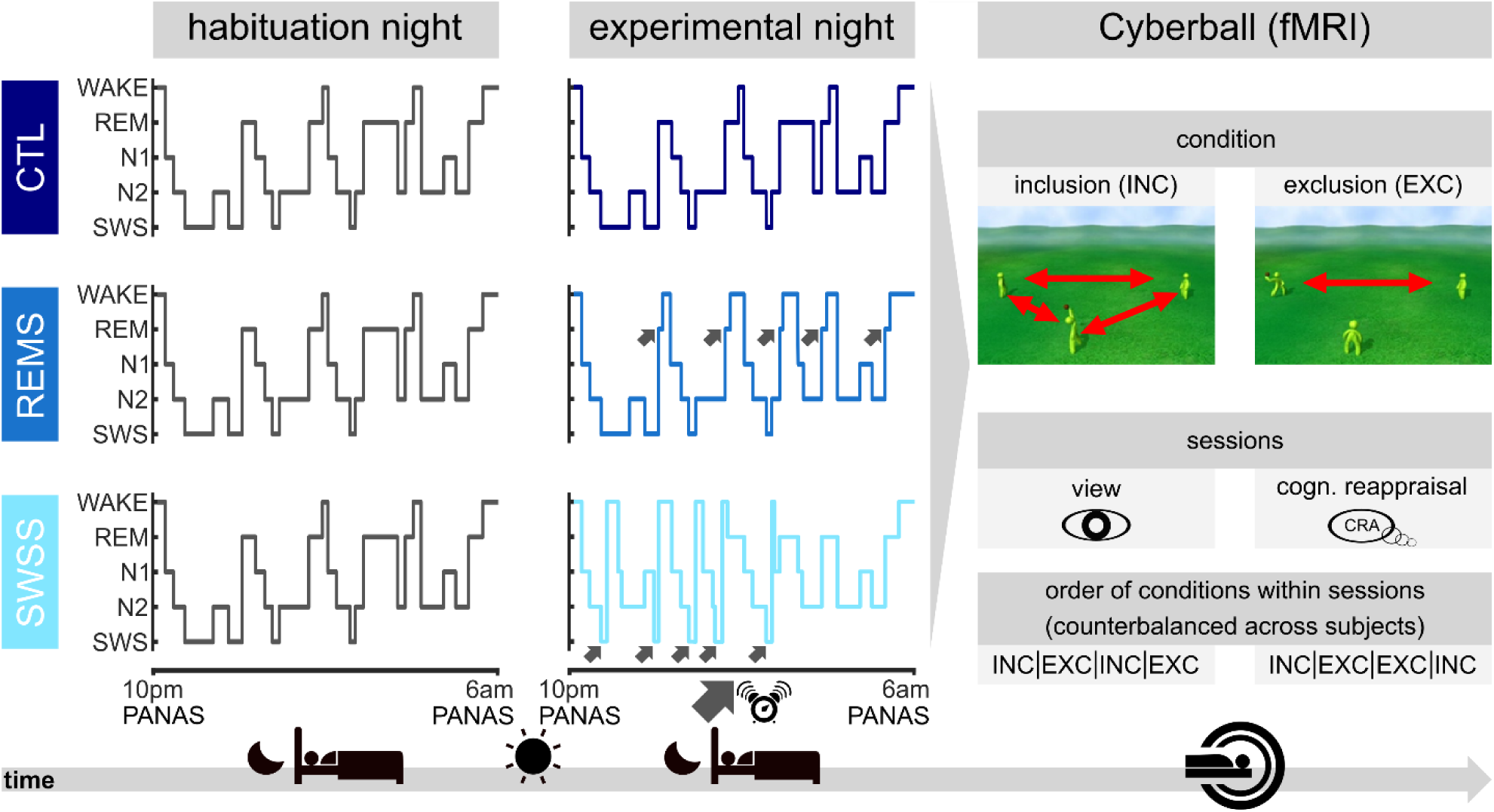
Summary of the experimental procedure. Three groups of subjects spent two consecutive nights in the sleep laboratory. During both nights, polysomnography was recorded. In the second night, two groups were woken up as soon as they entered REM sleep or SWS (groups REMS and SWSS, respectively), while the control group was not woken up (CTL). After waking up on the second morning, all subjects performed two sessions of the *Cyberball* game while inside the fMRI scanner ^[92]^. The game included a total of eight blocks, with four inclusion (INC) and four exclusion (EXC) blocks. During inclusion, subjects received the ball continuously throughout the block, while during exclusion, the other two players stopped throwing the ball to the participant soon after the beginning of the block, effectively excluding the participant from the game. Additionally, during the first four blocks (half INC, half EXC), subjects simply performed the task (VIEW), while during the second four blocks (half INC, half EXC) they were instructed to use cognitive reappraisal (CRA) to regulate their emotions.

**Figure 2.**
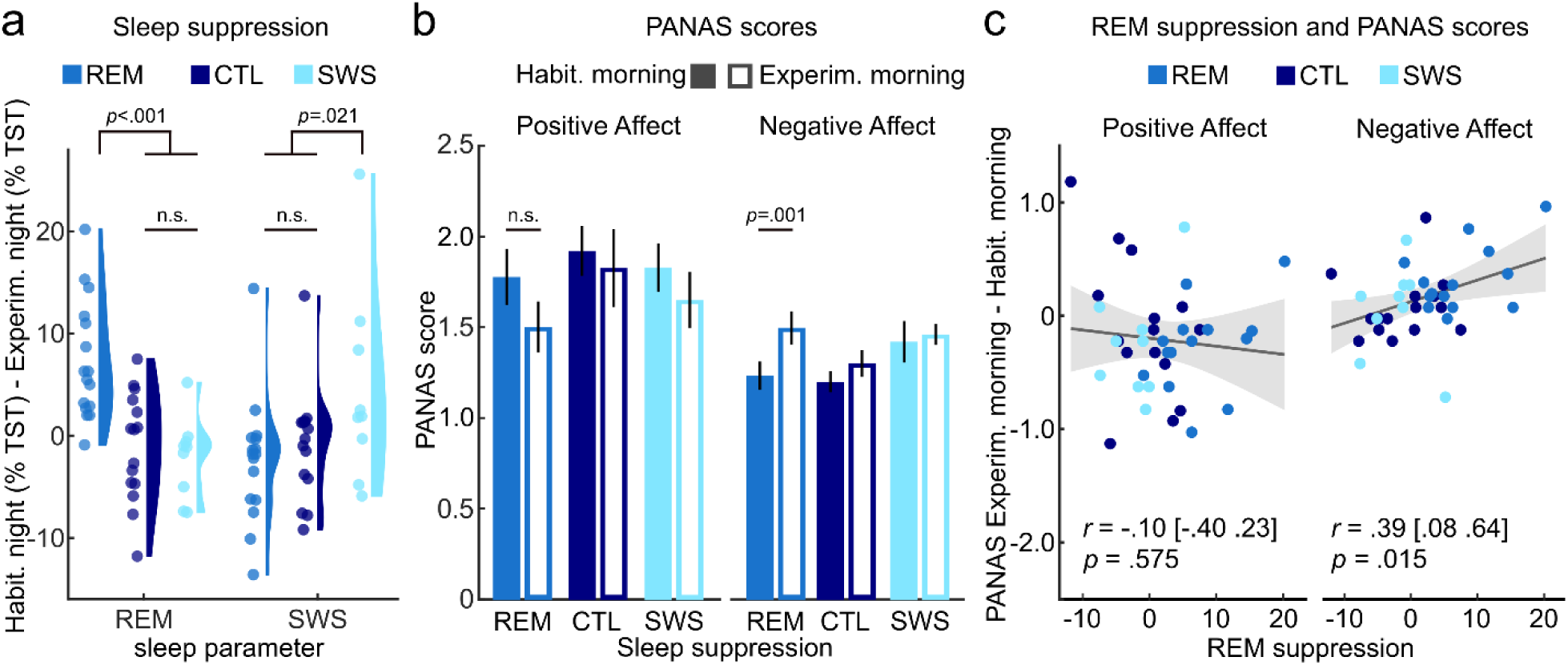
Selective sleep suppression and general affect. **a** REM sleep was significantly more reduced in the REMS group than in the other two groups, whereas SWS suppression was significantly stronger in the SWSS group than in the other two groups. Suppression scores for REM (SWS) sleep are the difference of percentage REM (SWS) of TST between the habituation night minus the experimental night. **b** Ratings of general positive and negative affect, measured with the Positive And Negative Affect Scale (PANAS)^[50]^ on the mornings after the habituation night and after the experimental night, separately for the three groups. **c** Changes in negative (right) but not positive affect (left) from the habituation night to the experimental night correlate significantly with REM sleep suppression scores across all groups. Shaded areas represent 95% confidence intervals for simple bivariate regression. Error bars represent standard error of the mean. n.s.=not significant.

### REM Sleep Predicts General Negative Affect

General affect ratings in the PANAS obtained in the morning differed significantly between habituation and experimental night (effect of night: positive affect (PA): *F*(1,37)=4.55, *p*=.040,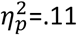; negative affect (NA): *F*(1,37)=9.33, *p*=.004, 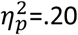), while type of suppression had no significant effect (PA: *F*(2,37)=0.74, *p*=.484, 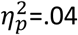; NA: *F*(2,37)=2.21, *p*=.124, 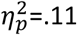). The interaction effect was not significant for positive affect (PA: *F*(2,37)=0.34, *p*=.713, 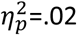) but approached significance for negative affect (NA: *F*(2,37)=3.08, *p*=.058, 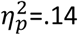). Across groups, post-hoc comparisons demonstrated that positive affect was lower and negative affect was higher after the experimental night as compared to the habituation night (PA: *t*(39)=2.20, two-sided *p*=.034, *d*=0.35, 95% *CI*=[0.03;0.67]; NA: (*t*(39)=-3.23, two-sided *p*=.003, Cohen’s *d*=-0.51, 95% *CI*=[-0.84;-0.18]), with the increase in negative affect being most pronounced in the REMS group (*t*(14)=4.47, *p*=.001, Cohen’s *d*=1.15, 95% *CI*=[0.48;1.80]; see figure 2). A planned contrast showed that this increase in negative affect was stronger in the REMS group as compared to the other groups (*t*(38)=2.47, two-sided *p*=.037, Cohen’s *d*=.81, 95% *CI*=[0.14;1.47]), whereas no group difference could be observed for change in positive affect (*t*(38)=-0.72, two-sided *p*=.950, Cohen’s *d*=.24, 95% *CI*=[-0.88;0.41], all *p*-values Bonferroni-corrected). Regarding the evening ratings, type of sleep suppression did not have any significant effects or interactions, neither for PA nor for NA (all *p*s>.110; see supplementary information for details). Across groups, increases in morning negative affect from habituation to experimental night could be predicted by the REMS score (Pearson’s *r*=.39, one-sided *p*=.015, 95% *CI*=[0.08;0.63]; figure 2), even after controlling for changes in SWS, total sleep time (TST) and wakefulness after sleep onset in a multiple linear regression model (see table 2).

**Table 2.**
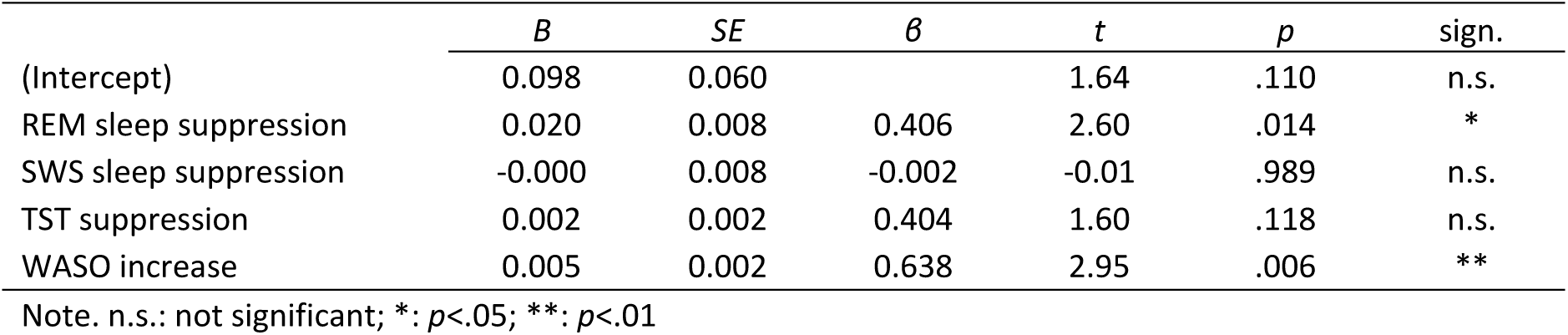
Regression of affective changes on sleep deprivation parameters (dependent variable: change in negative affect (Δ Experimental morning – Habituation morning))

| | *B* | *SE* | $\beta$ | *t* | *p* | sign. |
| --- | --- | --- | --- | --- | --- | --- |
| (Intercept) | 0.098 | 0.060 |  | 1.64 | .110 | n.s. |
| REM sleep suppression | 0.020 | 0.008 | 0.406 | 2.60 | .014 | * |
| SWS sleep suppression | -0.000 | 0.008 | -0.002 | -0.01 | .989 | n.s. |
| TST suppression | 0.002 | 0.002 | 0.404 | 1.60 | .118 | n.s. |
| WASO increase | 0.005 | 0.002 | 0.638 | 2.95 | .006 | ** |

Note. n.s.: not significant; \*: $p < .05$ ; \*\*: $p < .01$

### No Effect of Selective REM Sleep Suppression on Emotional Responses to Social Exclusion

By means of the *Cyberball*, we successfully induced the unpleasant feeling of social exclusion (main effect of INC/EXC: *F(*1,39)=131.65, *p*<.001, 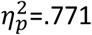; see table 3 for descriptive statistics), thereby replicating earlier findings ^[48]^. Moreover, if participants were asked to engage in emotion regulation using cognitive reappraisal, feelings of being excluded could be toned down significantly (main effect of VIEW/CRA: *F(*1,39)=66.99, *p*<.001, 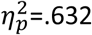, with this effect being most strongly pronounced during trials of social exclusion (interaction effect INC/EXC x VIEW/CRA: *F(*1,39)=44.97, *p*<.001, 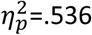).

**Table 3.** Feelings of being excluded, separately for groups and conditions

|  | EXC/VIEW |  | INC/VIEW |  | EXC/CRA |  | INC/CRA |  |
| --- | --- | --- | --- | --- | --- | --- | --- | --- |
|  | *M* | *SD* | *M* | *SD* | *M* | *SD* | *M* | *SD* |
| CTL | 6.67 | 1.92 | 2.20 | 1.25 | 3.67 | 2.50 | 1.50 | 0.78 |
| REMS | 6.38 | 2.90 | 2.00 | 1.21 | 2.82 | 2.33 | 1.62 | 0.86 |
| SWSS | 5.90 | 2.49 | 2.25 | 0.98 | 2.60 | 1.51 | 1.75 | 1.46 |

However, regarding our second hypothesis, type of sleep suppression neither had a significant impact on emotions after social exclusion (INC/EXC * group interaction: *F(*2,39)=1.47, *p*=.243, 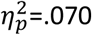) nor on the effect of emotion regulation (VIEW/CRA * group interaction: *F(*2,39)=.027, *p*=.973, 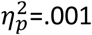). Last, we did not find a statistically significant three-way interaction on the self-reported emotions after social exclusion (INC/EXC * VIEW/CRA * group interaction: *F(*2,39)=.45, *p*=.639, 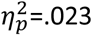). Similarly, no significant main effect of or interactions with type of sleep suppression were found when contrasting REMS against the other two groups (all *p*-values >. 41).

### REM Sleep Suppression Alters Amygdala Activity during Social Exclusion

Similar to earlier studies, we found that during social exclusion participants’ neural activity in the left and right hippocampus (left: x,y,z (mm): -36,-40,-6; *T*=7.15, *k*=89, *p*<.001, FWE-corrected; right: 36,-32,-8; *T*=4.88, *k*=4, *p*=.022, FWE-corrected) as well as the right insula (34,-10,22; *T*=5.07, *k*=13, *p*=.014, FWE-corrected) was significantly increased compared to the inclusion condition ^[51,52]^. In addition, across groups, neural activity in the right anterior insula was significantly increased during VIEW as compared to CRA blocks (main effect VIEW>CRA: 30,24,-4; *T*=4.58, *k*=1, *p*=.048, FWE-corrected). Testing whether these two main effects interacted or whether they were modulated by type of sleep suppression did not yield any significant effects. This held both when testing for differences between any of the three experimental groups and when comparing the REMS group against the other two groups combined.

Last, we tested the three-way interaction of EXC/INC, VIEW/CRA, and type of sleep suppression. A test of differences between any of the three groups for the two-way interaction contrast [EXC/VIEW>INC/VIEW] > [EXC/CRA>INC/CRA] was not significant. However, comparing the REMS group to the other two groups indicated differential neural responses in the right amygdala for the REMS group (26, -2, -28; *F*=23.70, *k*=1, *p*=.048, FWE-corrected inside the a priori mask; see figure 3). Precisely, the contrast EXC>INC was positive during VIEW in the REMS group, but did not differ from zero in the other groups (REMS: *t*=4.04, two-sided *p*=.005; CTL: *t*=-1.07, two-sided *p*=1.000; SWSS: *t*=-0.58, two-sided *p*=1.000; see figure 3). In addition, EXC>INC was more positive during VIEW than during CRA in the REM group (*t*=5.27, two-sided *p*<.001), but not in the other groups (CTL: *t*=-2.18, two-sided *p*=.141; SWSS: *t*=-1.70, two-sided *p*=.369; all p-values Bonferroni-corrected). This pattern of results suggests that the signaling of information that is relevant to the individual’s social well-being and mediated by amygdala activity depends on intact REM-sleep.

**Figure 3.**
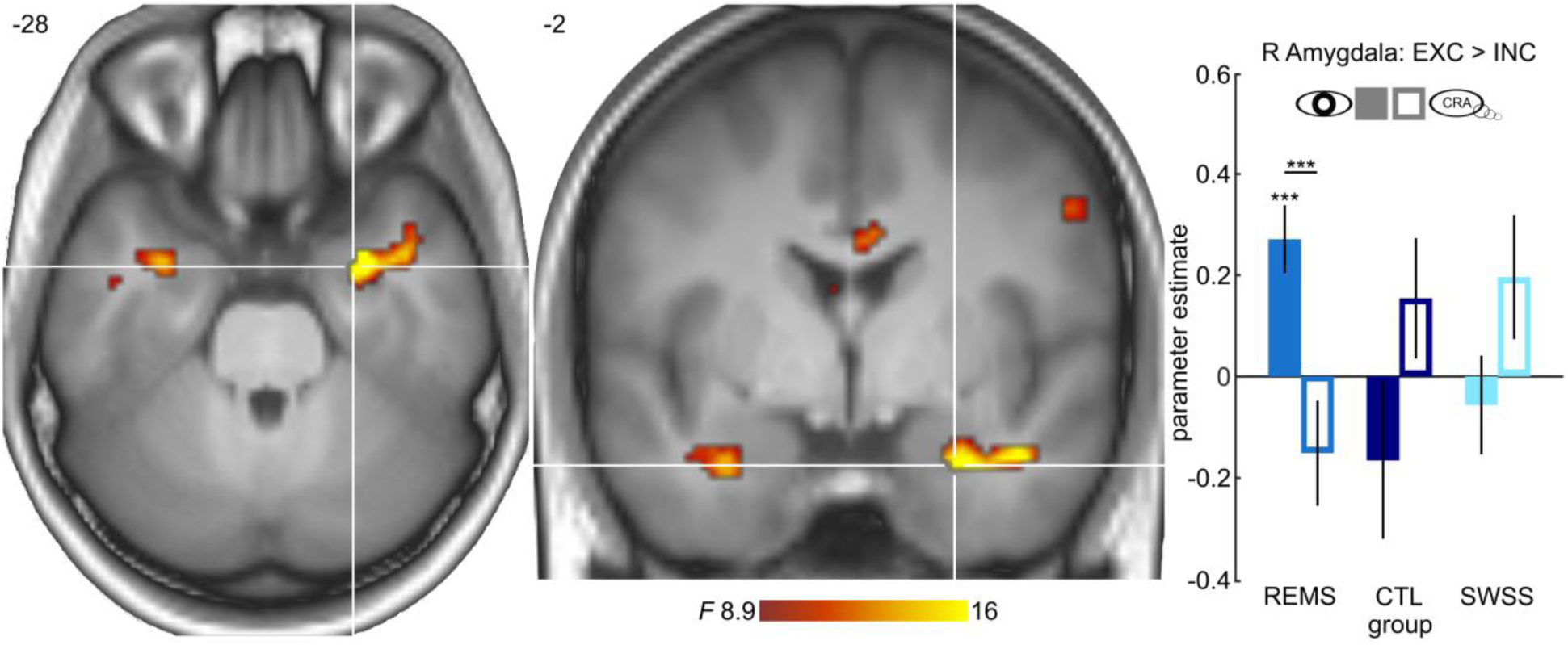
Neural responses to ostracism are altered by selective REM sleep suppression. **Left** Neural responses for the contrast [EXC/VIEW>INC/VIEW] > [EXC/CRA>INC/CRA] differed significantly between the REMS group and the other two groups in the right amygdala, (peak MNI-coordinates: x=26, y=-2, z=-28, surviving FWE-correction at *p*<.05 inside the a priori mask). Displayed at *p*<.005, uncorrected, for visualization purposes. **Right** Parameter estimates for the contrast EXC>INC in the VIEW session were positive in the REMS group, but did not deviate from 0 in the other two groups, nor in the CRA session for any of the three groups. In addition, parameter estimates for EXC>INC were significantly more positive in the REMS group during VIEW than during CRA, while sessions did not differ in the other two groups. See main text for details. ***: *p*<.001. Error bars represent standard error of the mean.

## Discussion

Nearly everyone can relate to the devastating effects of a sleepless or interrupted night on one’s next day mood. While the effect of total sleep deprivation on emotional reactivity has been investigated intensively in the past ^[53,54]^ the present study focused on the specific impact of selective REM sleep suppression on general affect, as well as emotion regulation and its neural correlates under conditions of social exclusion.

We found that lower amounts of REM sleep across all participants were associated with higher levels of general negative affect in the next morning, a finding that is in line with previous literature implicating REM sleep in emotional functioning ^[55]^. Despite this general effect, however, our findings do not provide evidence for a direct link of REMS with the subjective emotional response to experimentally induced social exclusion. The ability to regulate one’s negative emotions during social exclusion was also not affected by prior REMS, which was an unexpected finding. Interestingly though, despite no changes in subjectively reported emotions, neural activity in the limbic system was altered after REMS when participants experienced social exclusion. Precisely, neural responses to passively experienced ostracism were specifically increased in right amygdala after REM sleep was selectively suppressed. The amygdala is strongly associated with emotional processing ^[56,57]^ and has anatomical connections to the anterior insula and the anterior cingulate cortex ^[58,59]^. As part of this so-called salience network ^[60]^, the amygdala’s assumed function of signaling the relevance of information is central for the domain of affective experiences ^[57,61]^. Previous studies showed that amygdala responses to viewing negative emotional stimuli increased after total sleep deprivation ^[62]^, depended on intact REM sleep in particular ^[33]^, and correlated with autonomic responses to psychosocial stress ^[63]^. Our study connects these previous findings by demonstrating that amygdala activity tracks information relevant to the subjects’ social well-being in dependence of REM sleep.

We can only speculate why these alterations in brain function after REMS did not manifest on the behavioral level in form of increased emotional reactivity to the experience of social exclusion. Previous research provides several explanations that might be helpful to understand this dissociation. First, the effects of experimental short-term sleep manipulations might be strongest immediately after awakening and may be readily washed out thereafter ^[64]^, and may thus not have lasted until the fMRI session in the present study. This may hold in particular for selective suppression of specific sleep stages rather than total sleep deprivation, which produces stronger and more long-lasting effects ^[64]^. The efficacy of REMS in the present study may have been too small to surface on the behavioral level later in the morning. However, the effects of REMS nonetheless persisted on the level of brain systems. The altered brain activity in the amygdala could indicate that even small changes in REM sleep can influence brain systems implicated in regulating responses to affectively salient stimuli ^[56,57]^, while they are too small to penetrate the level of subjective experiences, which are possibly processed further downstream ^[63,65]^. Yet, what speaks against this explanation are findings by Wiesner and colleagues ^[30]^ or Morgenthaler and colleagues ^[66]^ who applied even more rigorous REM sleep deprivation, achieving a mean REM sleep percentage of around one percent of TST, but nevertheless could not find REM sleep related behavioral effects during emotion recognition tasks. Similarly, Liu and colleagues compared the effect of a 24h sleep deprivation to regular sleep on the experience of distress following social exclusion in the *Cyberball* game ^[67]^. As in the present study, the authors did not find an effect of sleep deprivation on the subjective experience of social rejection.

Regardless of the timing and potential washout during the day, the lack of significant findings for the experience of social exclusion might also relate to the conceptual breadth and the ecological validity of the affective assessment. While the underlying brain systems show increased reactivity towards potentially threatening conditions and signal greater homeostatic imbalance, the distinct assessment of emotional responses to ostracism, such as feelings of exclusion, might touch a different facet of affective construal. That is, the affective salience of information tracked by amygdala activity ^[57,61]^ may be modulated by REMS. However, the construal of emotional experience is assumed not to rely on the activity in single regions, but on the dynamic interaction of various neural systems supporting multiple psychological processes of emotional experience apart from affective salience ^[68,69]^. Hence, amygdala responses to psychosocial stress do not necessarily influence the construal of subjective emotion ratings, that may depend on additional networks involving prefrontal cortical regions, as suggested by previous studies ^[63,70,71]^. In addition, we did not find evidence that REM sleep deprivation modulated the ability to apply cognitive reappraisal on the behavioral level. Therefore, REM sleep deprived subjects may have been able to counteract the impact of increased amygdala responses during social exclusion. A potential factor diminishing the severity of experimental social exclusion is that despite our attempts to create a socially immersive context ^[72]^, laboratory studies per se have limited relevance beyond the experimental situation itself. Thus, even without explicit reappraisal strategies, this limited relevance might have entered into the construal of emotional responses during passive social exclusion, and reduced the influence of altered amygdala responses in the REMS group.

A related explanation for the brain-behavior dissociation is that healthy participants have resources for compensation. As the neuroimaging findings advocate that REMS impacts limbic circuit activity during emotional experiences, the question arises whether participants with a priori (sub)clinical peculiarities in reaction to social exclusion, ostracism or labile sense of belonging would also report altered subjective experiences. It has been shown that inter-individual dispositions such as differences in rejection sensitivity ^[73]^, social anxiety ^[74]^, trait self-esteem, depression ^[75]^, attachment style ^[76]^ and situational factors of the exclusion experience moderate the effect of social exclusion. Last, it is possible that experimental REM-sleep suppression becomes effective on the emotional level only when applied repeatedly, simulating chronic sleep disturbances that are associated with psychiatric disorders in a more realistic manner.

Speculatively, a further explanation could relate to the psychological task used to examine emotional reactivity. Earlier research indicated that sleep deprivation effects might differ depending on whether the focus was on general state-like morning affect (e.g. as measured using the PANAS), the processing and responding to affective material (e.g. emotion recognition, pain processing, reactivity to threatening stimuli) or on the direct induction of emotional states (e.g. inducing the unpleasant experience of social exclusion). One assumption derived from the present data could be that direct emotion-induction tasks are so powerful that a single night of sleep deprivation or selective REMS might not suffice to exert effects on the psychological level. To our knowledge, however, apart from the present study there is no further work that directly examined the effect of selective REMS on experimentally induced emotional states. Second, general unspecific affect may function differently than emotional reactions to specific elicitors ^[77]^, potentially moderating the effect of REMS on these different affective processes. However, at least one study found that general affect was not influenced by REMS, which contradicts our findings ^[30]^. Last, we are not aware of any study systematically comparing the extent to which the different psychological aspects of affective experience are susceptible to selective suppression of sleep stages. Taken together, the specific interaction of selective REMS with general affect, in contrast to more confined emotional responses, demands more in-depths analyses of different kinds of experimental designs and dependent variables.

Our findings of increased next morning general negative affect as well as increased limbic activity after REMS are well in line with findings in patients with mental disorders. A wide range of mental disorders, including mood and anxiety disorders, are not only accompanied by profound disturbances of REM sleep ^[78,79]^ but also by deficits in generating and controlling emotions in an appropriate way. Posttraumatic Stress Disorder (PTSD) is one of these mental disorders, where a link between REM sleep and emotion regulation has been frequently discussed ^[80,81]^. In these patients, deficits in emotion processing and regulation on the behavioral level are coincided by alterations in brain networks implicated in emotion regulation such as the amygdala, hippocampus, insula, and anterior cingulate ^[82]^. Apart from that, PTSD patients as compared to traumatized individuals who did not develop a PTSD, show a profound REM sleep fragmentation and alterations in REM density accompanied by higher sympathetic drive during this sleep stage ^[78,79]^. Importantly, these REM sleep disturbances early after trauma predict later PTSD development ^[83,84]^, which let researchers speculate that REM sleep alterations are not only a secondary symptom but represent one mechanistic factor contributing to the development and/or the exacerbation of PTSD symptoms ^[80,81]^. However, empirical evidence for such a mechanistic link is scarce.

Our findings of altered negative affect and changed brain responses to social exclusion after one night of REMS in healthy subjects resemble patterns typically seen in PTSD patients. Hence, the present data provide additional evidence for a role of REM sleep in emotion processing and regulation and they further suggest a mechanistic link between REM sleep disturbances and psychopathology. Future research should investigate the effect of experimental REM sleep manipulation (i.e. REM sleep increase as well deprivation/fragmentation) through pharmacological and/or psychological treatments (i.e. cognitive behavioral therapy for insomnia, CBT-I^[85,86]^) on emotion regulation in healthy subjects and patients suffering from mental disorders. Conducting more research in this field gains additional importance from the fact that sleep deprivation does not have negative effects only, but can also have positive effects, as evident in its anti-depressant potentials ^[87]^.

## Limitations

Although in the present study, the use of cognitive reappraisal (CRA) strategies was effective in toning down feelings of social exclusion ^[88]^, CRA was always applied in the second session. Therefore, we cannot rule out that habituation to the task during the second session influenced the regulation of emotionality by CRA. However, counterbalancing the view and reappraisal session would have introduced even stronger confounds, considering that participants would have most likely also engaged in CRA in the second session if they had learned about this strategy in the first session. In order to better discriminate between CRA and habituation effects one could apply a three-session-design and randomly instruct participants to use CRA either in the second or third session. In a larger sample than ours, Mauss and colleagues applied such a design to show that poorer sleep quality over the course of the last week was linked to decreased abilities in engaging in CRA ^[41]^.

A second limitation may be the presence of social desirability effects. While CRA is regarded as the most efficient emotion regulation technique ^[89]^, the experimenters’ intention may have been too obvious in the present social exclusion context, which might have increased social desirability effects, rendering this emotion regulation strategy suboptimal. Alternatively, in a different experimental setup, one could implicitly offer subjects an opportunity for diverting their attention away from the unpleasant emotional experience in one session and thereby apply another emotion regulation technique (i.e. suppression) without an explicit instruction and the danger of social desirability effects. Future studies will need to specifically target the effect of different emotion regulation strategies and the impact that selective REMS might exert on such techniques.

## Conclusion

In conclusion, the present study examined the effect of selective REMS on general affect as well as the neural correlates of task-induced negative emotionality and the ability to regulate emotional experiences by cognitive reappraisal during experimentally induced social exclusion. While we found that REMS predicted general negative affect in the next morning, we did not find experimental evidence for task-induced changes of negative emotionality during the experience of social exclusion. However, limbic system activity during social exclusion was significantly increased in REM sleep suppressed participants pointing to a possible neural mechanism linking REM sleep disturbances and affect. Further investigations of the underlying mechanisms of REM sleep and emotional dysregulation in social situations will pave the way for a better understanding of the role of sleep disturbance in psychiatric disorders.

## Methods

### Participants

A total of 45 participants were initially invited to take part in the experiment (29 female, age (years): *M*=23.69, *SD*=2.67), gave written informed consent prior to participation in the study and received financial compensation for their participation. All participants were recruited at Philipps-University Marburg, were fluent in German, and had normal or corrected-to-normal vision. None of them were diagnosed with neurological or psychiatric disorders (present and past), current alcohol or drug abuse, use of psychiatric medications (present and past), anatomical brain abnormalities (e.g. lesions, strokes etc.), or sleep disturbances. Due to technical issues with the polysomnographic recording in the experimental night (n=1) or insufficient quality of the fMRI data (n=2), 3 subjects were not included in the final analyses. The final sample thus consisted of 42 participants (27 female, age (years): *M*=23.76, *SD*=2.74). Participants were randomly assigned to one of three groups, which differed regarding to the sleep protocol in the experimental night (i.e. either REMS, SWSS or CTL, details below). Groups did not differ in age (CTL: *M*=24.00 years, *SD*=3.12; REMS: *M*=23.06, *SD*=2.22; SWSS: *M*=24.60, *SD*=2.91; *F*(2,39)=1.09, *p*=.346) or gender (CTL: 8 female, 7 male; REMS: 12 female, 5 male; SWSS: 7 female, 3 male; *X*^2^(2,42)=1.22, *p*=.543).

### Sleep Manipulation Procedure

Participants came to the laboratory for two consecutive nights – one habituation night and a subsequent experimental night (see figure 1). The purpose of the habituation night was to exclude sleep disorders, make participants familiar with the polysomnography (PSG) recording procedure and adapt to the environment in the sleep laboratory. The morning after the experimental night, participants were taken to the fMRI facility where they completed the experimental task in the scanner (see Experimental task below). During both nights PSG was recorded to monitor and identify sleep phases (see table 1). A maximum of three participants was invited to the sleep laboratory on each night. Upon arrival on the habituation night, participants were informed about the procedure of the study and gave written informed consent. After the habituation night participants could spend the day as usual but were asked to refrain from napping, smoking and consuming stimulating foods and drinks (e.g. coffee, tea, energy drinks). On the subsequent night, participants entered a fully equipped single sleep room for PSG (Embla N7000 PSG, TNI-Medical, Würzburg, Germany) consisting of electroencephalography (EEG), electrooculogram (EOG), electromyogram of the mentalis muscle (EMG) and pulse oximetry using EMBLA N7000 (TNI-Medical). The EEG system had 9 electrodes positioned according to the 10-20 system, adhering to the guidelines of the American Academy Of Sleep Medicine (AASM)^[90]^, and placed on the following positions: F3, F4, C3, Cz, C4, O1, O2, M1 and M2. Limb movements were videotaped with infrared camera. The experimenter turned off the lights at 10 pm and asked participants to try to fall asleep. In case a deterioration of the PSG recordings was observed, the experimenter entered the sleep room to improve the signal by reattaching the respective electrodes. At 6 am, lights were turned on, participants were woken up and the electrodes were removed.

For those participants in the REMS and SWSS group, during the second night (i.e. the experimental night), the experimenter started an acoustic beep (80 dB, 500 Hz, 500ms) ^[91]^ once the polysomnographic trajectories indicated that the participant had entered the target sleep phase. After an awakening, participants were kept awake for 90 seconds. In case the participant did not wake up, the volume of the acoustic beep was increased, and ultimately the experimenter entered the participant’s room to turn on the lights to make sure the target sleep phase was interrupted. To control for the number and lengths of awakenings, both suppression groups were disturbed similarly often during the experimental night. In the REMS group, participants’ sleep was disturbed as soon as they entered the REM sleep stage. Awakenings for the SWSS group were selectively carried out during the non-REM sleep stage N3, defined according to AASM guidelines ^[90]^. The control subjects were not awakened at all. PSG recordings were scored by an experienced sleep technician (C.L.) at the sleep laboratory at the Department of Otorhinolaryngology at the University of Lübeck according to AASM guidelines ^[90]^ and using Somnologica 3.3.1 (Build 1529). The technician was blinded for group assignments and hypotheses.

### General Affect Ratings

The PANAS ^[50]^ was completed shortly before going to bed and after waking up on both nights by all participants to assess the effect of sleep manipulation on general positive and negative affect.

### Experimental Task

Upon waking up after the experimental night, participants were accompanied to the functional magnetic resonance imaging (fMRI) facility. In the MRI, participants were instructed via a screen that they were about to play a ball-tossing game, i.e. the *Cyberball* task, allegedly with two other persons (see Figure 1; a second paradigm was presented to the subjects afterwards, which involved IAPS pictures to study regulation of basic emotions but is not further described in the present manuscript). All three players, including the participant, were represented by avatars, i.e. green, 3D-animated stick-figures standing in a triangle on a lawn (for a detailed description of the animation see ^[92]^). Participants were able to control the avatar on the bottom edge of the screen, while the other two avatars were placed on the left and right of the horizontal midline of the screen. The names of the participant and of the other two players were displayed next to the respective avatar. For each throw, a short sentence on the bottom of the screen indicated who threw the ball to whom. When one of the computer avatars had the ball, they waited a random amount of time between 1000 and 2000 ms before tossing the ball. When the participant had the ball, they had 2000 ms to decide where to toss the ball. In case they responded too slowly, the computer randomly selected which of the other avatars would receive the ball. The duration of each throw was 750 ms and was visualized by a series of 40 frames, showing an avatar doing a throwing movement and the red ball flying from one avatar to the catching avatar, with the latter moving its arm to catch the ball. While participants believed that the other avatars were controlled by two other anonymous persons sitting in adjacent rooms, their behavior in fact followed a predefined script (see below). Following the instructions, a short sequence of lines of text built up on the screen, making the subjects believe that the computer was being connected to a gaming server of the university on which the experiment was run. This included the request of entering an IP-address, as well as lines stating how many players were logged into the game. Subsequently, participants performed a short training session consisting of seven trials in which they received and threw the ball three times.

The task then consisted of two sessions, each of which consisted of four experimental blocks. Before each of the experimental blocks, a line of text was presented on-screen for 1500 ms telling the participant that a new round would start. In every block, a maximum number of 24 ball tosses were completed. In two blocks of every session, the participant was included in the game and repeatedly received the ball throughout the entire block (inclusion blocks, INC), having the possibility to toss the ball to one of the other avatars eight or nine times by clicking a button using the index finger (left player) or middle finger (right player). In the other two blocks (exclusion blocks, EXC), after the participant had received the ball three or four times, the paradigm was programmed so that the two avatars only tossed the ball between one another, thereby effectively excluding the participant from the game.

After having completed four blocks during which subjects simply participated in the task without additional instructions (VIEW session), a second session of the task was played. For the second session, a short on-screen text instructed participants to use cognitive reappraisal to regulate their emotions during the game (CRA session; see Fig. 1). Participants were specifically asked to reappraise the situation, in case any negative emotions should arise (“In case negative emotions arise, please try to re-evaluate the situation”). This was accompanied by the instruction to “partake in the game and try to visualize the situation as vividly as possible”, which was also presented before the first session. Subjects were asked to press the left button as soon as they were ready, and then a short break varying randomly between 1500 and 2500 ms was presented, instructing participants to wait until the other two players had indicated that they were ready. After presentation of a fixation cross for one second, the block started.

In each session, two inclusion and two exclusion blocks were presented to the subjects, with the first block of each session always being an inclusion block. In one session, inclusion and exclusion blocks were alternating, while in the other session, the initial inclusion block was followed by two consecutive exclusion blocks and a final inclusion block. The assignment of block order to the VIEW and the CRA sessions was counterbalanced across subjects. On average, EXC blocks lasted 50.93 seconds (*SD*=0.62), and INC blocks lasted 45.87 seconds (*SD*=2.21).

Between blocks, participants were asked to rate how they felt about the preceding block regarding the extent to which they felt socially excluded. The low end of the scale was labelled as rejected/despised and the high end was labelled as accepted/familiar. In addition, subjects rated their sadness, anger, and shame, but these ratings were intended to distract from the purpose of the study and not analyzed (sadness was described by: sad, downcast, gloomy; for anger: angry, irritated, furious, mad; for shame: abashed, embarrassed). Each rating was displayed using a 9-point Likert-type scale ranging from 0 (not at all) to 8 (very much). Every rating was initialized at 4 (i.e. a neutral rating), and participants used presses of the right or left response buttons to move the rating to higher or lower values, respectively. The time for ratings was limited to 4 seconds. After each rating and the start of the next block within a session there was a short pause, jittered between 4.3 to 5.3 seconds. Emotion ratings for each condition (EXC, INC) and emotion regulation session (VIEW, CRA) were averaged for each participant and analyzed using repeated measures analyses of variance (rmANOVA).

### Statistical Analysis

Average ratings of feeling excluded for each condition (EXC, INC) and session (VIEW, CRA), as well as the percentages of the different sleep stages for the habituation and experimental nights were analyzed using analyses of variance (ANOVA). Significant interactions and main effects were followed up using paired comparisons. Since we expected that selective REMS would specifically alter affective experience and associated neural responses ^[22,23]^, we furthermore performed an a priori planned contrast of the REMS group against both control groups, the CTL and SWSS. The alpha-level was set to .05, and was adjusted using Bonferroni-correction, in case multiple tests were performed.

### FMRI Data Acquisition and Preprocessing

For each of the two experimental sessions, 130 functional volumes were recorded at 3T (Siemens Trio, Erlangen), of which the first three were discarded to allow for equilibration of T1 saturation effects. Functional volumes consisted of 36 ascending near-axial slices (voxel size=3*3*3 mm, 10% interslice gap, FOV=192 mm) and were recorded with TR=2200 ms, TE=30 ms, FA=90°. In addition, a high-resolution anatomical T1 image was recorded consisting of 176 slices (voxel size=1*1*1 mm, FOV=256 mm, TR=1900 ms, TE=2.52 ms, 9° FA).

The MRI data were analyzed using SPM12 in Matlab 2019b. Functional MRI images from each session of the *Cyberball* paradigm were slice-time corrected to the middle slice and spatially realigned. Subsequently, spatial normalization to MNI space was performed using unified segmentation ^[93]^ by estimating the forward deformation fields from the mean functional image and applying these to the realigned functional images. These spatially normalized images were then resliced to a voxel size of 2*2*2 mm and smoothed with an 8 mm full-width at half-maximum isotropic Gaussian kernel and high-pass filtered at 1/256 Hz.

### FMRI Data Analysis

The preprocessed functional images were statistically analyzed by a two-level mixed effects GLM procedure. For each participant, a statistical model was specified, including data from both experimental sessions (VIEW, CRA). For each session, the INC, EXC and rating phases were modelled as regressors of interest and the six realignment parameters estimated during spatial realignment were included to account for variance in the functional data that was due to head motion. The contrast images obtained from the individual participants were then aggregated in a random effects model on the second level to test for effects of condition and session, and to test for differences between the three experimental groups. To disentangle significant interaction effects, we ran a series of post-hoc tests on the parameter estimates extracted from the peak activation, applying Bonferroni-correction for multiple comparisons.

### Regions of Interest (ROI) for FMRI Analysis

In order to focus our analyses on the limbic system as well as the insula, regions commonly associated with affective processing ^[94]^, we constructed an a priori mask using the Wake Forest University Pickatlas (v. 2.4)^[95]^. This mask comprised the anterior cingulate cortex, bilateral insula, bilateral hippocampus, as well as left and right amygdala.

## Author Contributions

Prepared and conceived the study: W.C., K.K., U.K., S.K., S.W.; Collected the data: S.K., F.M.P., S.W; Analyzed the fMRI and behavioral data: R.G., F.M.P., L.M.P., S.K., D.S.S.; Analyzed the PSG data: A.S.; Interpreted the data and wrote the manuscript: S.B., S.D., R.G., S.K., F.M.P., L.M.P., D.S.S., I.W.

### Acknowledgements

We thank Konrad Whittaker and Tareq Naji for their help in conducting the polysomnography measurements and Christian Lange for support in scoring the PSG data.

## Data Availability

The datasets generated during and/or analyzed during the current study are available from the corresponding author on reasonable request.

## Notes

### Competing Interest Statement

The authors have declared no competing interest.

### Summary of Updates

Restructured the manuscript with methods section last. New referencing style. Corrected minor mistakes.

