## Supplementary_Information_REM_SocialExclusion for "Selective REM-Sleep Suppression Increases Next-Day Negative Affect and Amygdala Responses to Social Exclusion" (from another version of this preprint: version 1, 2020-06-16)

**Supplementary Table 1.** ANOVA of evening Positive Affect ratings (PANAS; Crawford & Henry, 2004)

| Within Subjects Effects | Sum of Squares | *df* | Mean Square | *F* | *p* |
|---|---|---|---|---|---|
| Evening | 1.60 | 1 | 1.60 | 9.72 | .004 |
| Evening * REMSgroup | 0.03 | 1 | 0.03 | 0.16 | .688 |
| Residual | 6.24 | 38 | 0.16 | | |
| Between Subjects Effects | | | | | |
| REMSgroup | 0.47 | 1 | 0.47 | 0.94 | .339 |
| Residual | 19.13 | 38 | 0.50 | | |

Note. The PANAS was scored in the evenings before the habituation night and the experimental night (factor Evening). The REMS group was compared against the other two groups (CTL, SWSS; factor REMSgroup).

**Supplementary Table 2.** ANOVA of evening Negative Affect ratings (PANAS; Crawford & Henry, 2004)

| Within Subjects Effects | Sum of Squares | *df* | Mean Square | *F* | *p* |
|---|---|---|---|---|---|
| Evening | 0.14 | 1 | 0.14 | 1.99 | .167 |
| Evening * REMSgroup | 0.19 | 1 | 0.19 | 2.68 | .110 |
| Residual | 2.65 | 38 | 0.07 | | |
| Between Subjects Effects | | | | | |
| REMSgroup | 0.02 | 1 | 0.02 | 0.12 | 0.732 |
| Residual | 7.82 | 38 | 0.21 | | |

Note. The PANAS was scored in the evenings before the habituation night and the experimental night (factor Evening). The REMS group was compared against the other two groups (CTL, SWSS; factor REMSgroup).
